# The Origin and Migration of the Ameru Community in Kenya based on mtDNA analysis

**DOI:** 10.64898/2026.04.16.718862

**Authors:** David Miruka Onyango, Raphael Anampiu, Cyrus Ayieko, Lilian Magonya, Roselida Achieng Owuor, Gordon Obote Magaga, Brian Andika

## Abstract

Human diversity did not only remain restricted to their socio-cultural and linguistic domains but also have penetrated deep inside their genetic root. Africa harbors more genetic diversity than any other part of the world. Diversification of the African lineages were complex, involving long-distance gene flow. Data from Africansis needed to better understand the origin and evolution of modern humans, the genetic basis local adaptation, and the evolution of complex traits and related diseases. This analysis formed the basis for this study of determining the origin and migration of the Ameru community in Kenya. Blood samples was collected from 132 male adults of 65 year and above. DNA was extracted and analyzed for the Hyper variable region 1and 2. The sequences were sequenced using Sanger sequence alignment and analyzed using Geneious. Phylogenetic analysis was done using Mega-X while haplotype analysis was done using DNASP software. L1 haplogroup (2.9%) was found among Igembe (7%), Tharaka (6%), and Chuka (7%) and is common in West, Central, and parts of East Africa. L2 haplogroup (6.7%) was present in all subgroups except Imenti and Tigania, indicating West and Central African maternal ancestry. L1 and L2 haplotypes indicate that most Ameru subgroups share partial maternal ancestry from West and Central Africa, while Imenti and Tigania have different maternal lineages. L0–L4 haplogroups indicate predominant East, Central, and West African maternal origins, with subgroups showing variation in haplotype frequencies (e.g., L1 and L2 in Igembe, Tharaka, Chuka; L3 in Tharaka, Mwimbi, Chuka; L4 across all subgroups). Subgroup differences suggest that certain communities, particularly Imenti, have distinct maternal lineages, with less contribution from L1, L2, and L3 but potential links to Afro-Asiatic groups via L4 (found in the Middle East). Non-African haplogroups (N and R) point to historical interactions or shared ancestry with populations in Eurasia and the Horn of Africa, primarily in Tigania and Imenti. Overally, the Ameru maternal gene pool is heterogeneous, shaped by multiple migration routes and interactions across East Africa and beyond, with subgroup-specific maternal histories.

## Introduction

The Ameru of Kenya are a Bantu-speaking community settled on the eastern and northeastern slopes of Mount Kenya and the surrounding regions. Linguistically classified within the Northeast Bantu group, they form part of the wider demographic processes associated with the Bantu expansions across sub-Saharan Africa. Although collectively identified as Ameru, the community is internally differentiated into several sub-groups—including the Imenti, Tigania, Igembe, Tharaka, Mwimbi, Muthambi, Chuka, and Igoji—each of which preserves distinct genealogical accounts of origin and settlement. These narratives, together with linguistic affiliation, situate the Ameru within the eastern African highland Bantu milieu while maintaining claims of differentiated descent.

Any investigation of Ameru origins must be framed within the broader context of African population history. Modern humans originated in Africa within the past ∼300,000 years and have maintained continuous occupation of the continent (Fan et al., 2019). Genetic studies consistently demonstrate that Africa harbours greater genetic diversity than any other region of the world (Cann et al., 1987; Tishkoff et al., 2009), and that lineages found outside Africa represent a subset of African diversity (Tishkoff et al., 2009). The depth and structure of African genetic variation reflect complex demographic processes, including long-distance gene flow and early lineage diversification. Consequently, African populations are central to understanding both global human origins and regionally specific histories of migration and interaction. Data from sub-Saharan Africa, in particular, remain indispensable for reconstructing prehistoric demographic events and clarifying patterns that are not easily resolved from non-African datasets alone (Fan et al., 2019).

Within this continental framework, Ameru oral traditions present divergent accounts of ultimate origin. One prominent strand locates an ancestral homeland at Mbwa (or Mmbwa), often described as a coastal or island setting inhabited by people portrayed as phenotypically distinct from present-day Ameru. The identity of these inhabitants is uncertain: they may reflect memories of Arab settlements along the Swahili coast, earlier Bantu-speaking populations, or other Indian Ocean communities. Other traditions assert a Middle Eastern origin, while still others associate Ameru beginnings with Meroë in the Nile Valley. These narratives raise substantive historical questions. If interpreted literally, they imply either extra-African ancestry or sustained interaction with Afro-Asiatic, Arabian, or Nile Valley populations prior to settlement in the Mount Kenya region. At the same time, their internal inconsistencies suggest the possibility of layered memory shaped by migration, incorporation, and reinterpretation over time.

Genetic analysis—particularly of mitochondrial DNA (mtDNA)—provides an independent line of evidence with which to evaluate these claims. MtDNA is inherited maternally and does not undergo recombination, enabling mutations to accumulate along discrete maternal lines (Achili et al., 2005). These mutations define haplotypes, which are grouped into haplogroups on the basis of shared ancestral variants. Haplogroups often exhibit geographically patterned distributions that reflect ancient dispersals and subsequent demographic events. As a result, mtDNA variation has been widely applied in anthropological genetics to infer maternal ancestry, reconstruct migration routes, and assess degrees of relatedness among populations. In African contexts, mtDNA has contributed substantially to debates on the Bantu expansions, regional continuity, and historical interaction between Bantu-speaking and Afro-Asiatic– speaking groups. (Steflova et al., 2011; Veeramah et al., 2010)

In the case of the Ameru, mitochondrial haplotype analysis offers a means of assessing whether traditions of coastal, Middle Eastern, or Nile Valley origin correspond to identifiable maternal lineages from those regions, or whether the community’s maternal ancestry is predominantly rooted in eastern Africa. By situating Ameru genetic data within the wider landscape of African diversity, this study seeks to clarify the historical relationships among Ameru sub-groups and to evaluate competing narratives of origin through a combined anthropological and genetic framework.

## Materials and Methods

This study was designed to investigate maternal lineage structure within the Ameru community in order to detect possible signatures of multiple migratory events. A cross-sectional sampling strategy was employed across nine Ameru dialect groups: Igembe, Tharaka, Igoji, Mwimbi, Tigania, Chuka, Imenti and Muthambi.

A total of 132 individuals were recruited. Eligibility criteria included self-identification as Ameru, affiliation with one of the nine dialect groups, and age ≥65 years. Older participants were preferentially selected to minimize the influence of recent demographic mobility and to increase the likelihood that sampled maternal lineages reflected earlier generational structure. Although mitochondrial DNA is maternally inherited, only male participants were enrolled in order to reduce redundancy arising from sampling close maternal relatives within the same household clusters.

All participants provided written informed consent prior to enrolment. The study objectives and procedures were explained in the relevant dialects through community engagement sessions. Focus Group Discussions (FGDs) and structured open-ended questionnaires were used to contextualize the genetic investigation within existing oral traditions and to document self-reported clan affiliation, sub-tribal identity, and perceived ancestral origins. These qualitative data were used to complement genetic analyses and to ensure accurate sub-group classification.

### Sample Collection

Peripheral venous blood samples were collected by a trained phlebotomist under standard aseptic conditions. Approximately 1 ml of whole blood was drawn into a sterile 2 ml EDTA vacutainer tube. Each sample was assigned a unique identification code corresponding to the participant’s dialect group and questionnaire record.

Immediately after collection, samples were stored in a temperature-controlled cool box maintained at approximately 8°C. Samples were transported to the laboratory within the same day for downstream molecular analysis (Figure 1).

**Figure 1.**
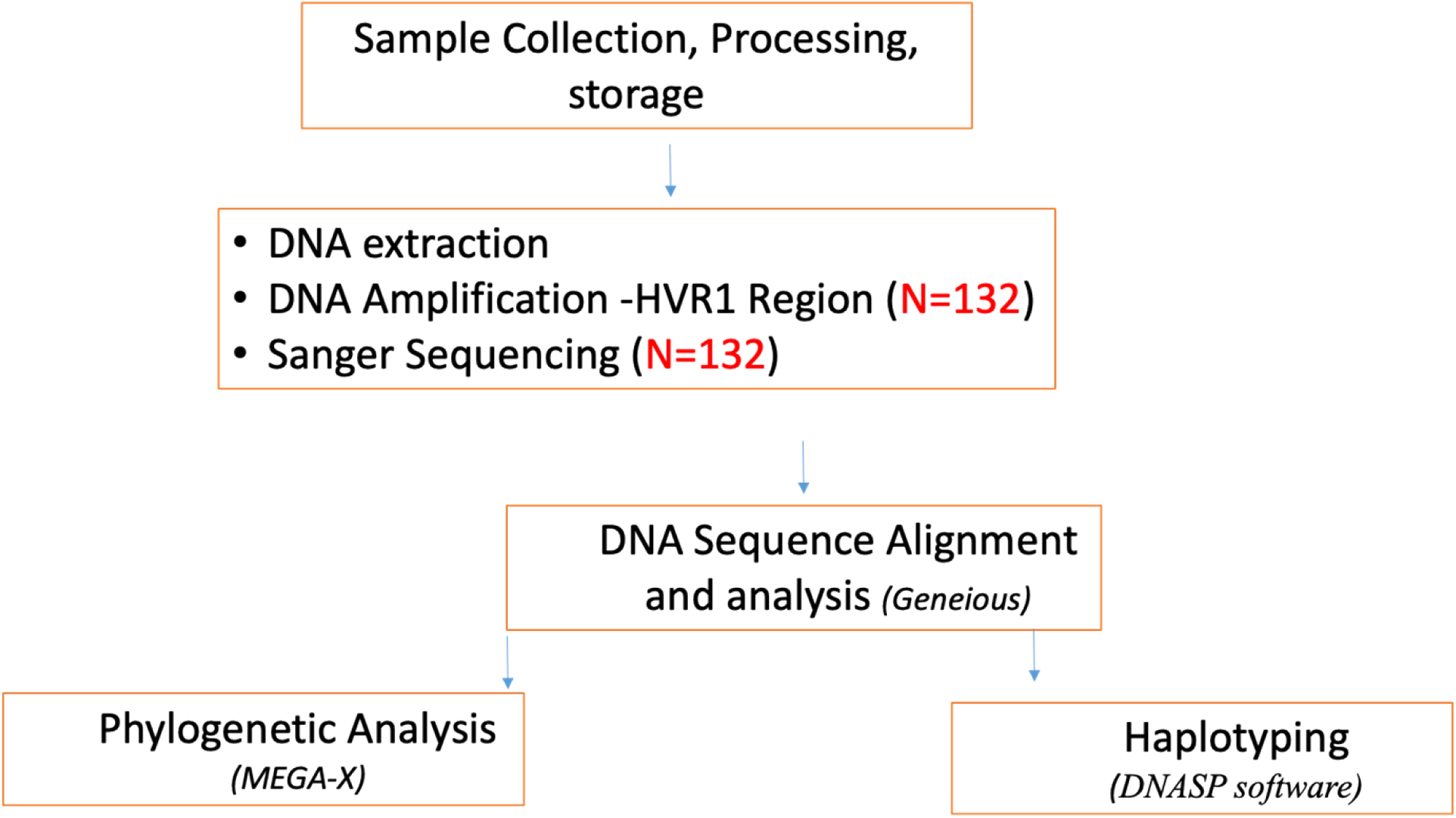
The flow chart and sequence of collected blood sample analysis.

### Sampling and sample preparation

The collected 132 human blood samples were then submitted to Sanger Labs at the Inqaba Biotec laboratories for DNA extraction and sequencing. Briefly, samples were aliquoted and 100ul of each sample in 200ul of DNA/RNA shield was extracted using the sample in DNA Shield protocol of the *Quick*-DNA Miniprep Plus Kit (Zymo Research, USA), DNA concentration and purity were quantified using the NanoDrop spectrophotometer, samples with a concentration above 20ng/ul and with nucleic acids purity ratios A260:A280 of 1.8 passed for PCR amplification.

### PCR Amplification of HVRI and HVRII

mtDNA from the previous step was amplified using hypervariable region primer sets HVRI-F(5’CTATCAACACACCCAAAGCTGAA3’) and HVRI-R (5’CGGAGCGAGAAGAGGGAT3’) spanning the loci 16024-16383, and HVR2-F (5’GGTCTATCACCCTATTAACCAC3’) and HVR2-R (5’CTGTTAAAAGTGCATACCGCCA3’) spanning loci 57-372. Briefly, PCR cycling conditions were as follows: Initial denaturation 94°C for 30 sec, denaturation at 94°C for 30 sec, annealing 53°C for 30 sec, elongation 68°C for 1 min and final elongation 68°C for 5 mins, using the One Taq Quick-Load 2X Master Mix (New England Biolabs, Massachusetts, USA) with a total reaction volume of 25μL. Amplicons were resolved on a 1.5-2% gel for HVRI and HVRII respectively.

### Sanger Sequencing D-loop region

Amplicons were enzymatically purified for any primer dimers using the EXO-SAP (New England Biolabs, Massachusetts, USA) with the following cycling conditions: step 1, 38°C for 15 mins, and step 2, 80°C for 15 mins. Purified amplicons were then cycle sequenced with HVRI and HVRII primer sets using the Brilliant Dye Terminator v3.1 (Nimagen, Netherlands) according to the manufacturers protocol.

### Bioinformatics analysis

Briefly, raw reads were parsed to CLC Main workbench v22.5 (Qiagen, Aarhus, Germany), data was trimmed and assembled using default settings. BLAST Searches were done to confirm the nucleotide sequences on the NCBI core database. Predicted haplogroup and variants were determined and analysed on the MTOMAP and MITOMASTER (Lott, M.T., et al. (2013)).

## Results

We sequenced mtDNA of 132 individuals from the Ameru ethnic community. Ancestral alleles were used to construct the phylogenetic relationship of the African Ameru and compared with Eurasia using neighbour-joining (NJ) method which assumed admixture. Thus, individuals who clustered near each other in the tree could either share a recent common ancestry and or experience gene flow. The boot strap value of most nodes are 1000. We identified 24 SNPs for HVR1, and 19 SNPs for HVR2, and selected a set of SNPs, after pruning based on linkage disequilibrium relationship of the Africa Ameru.

The haplotype analysis showed that upto 75% of the Ameru have L haplotype. This haplotype is shared with the rest of Africa populations. The predominant Haplotypes originate from East Africa i.e., L3(40%), L4(15%). The existence of haplotypes L0(20%), L1(3%), and L5(2%) evident of Southern and Central Africa haplotype is also reported (Fig.2)

**Figure 2.**
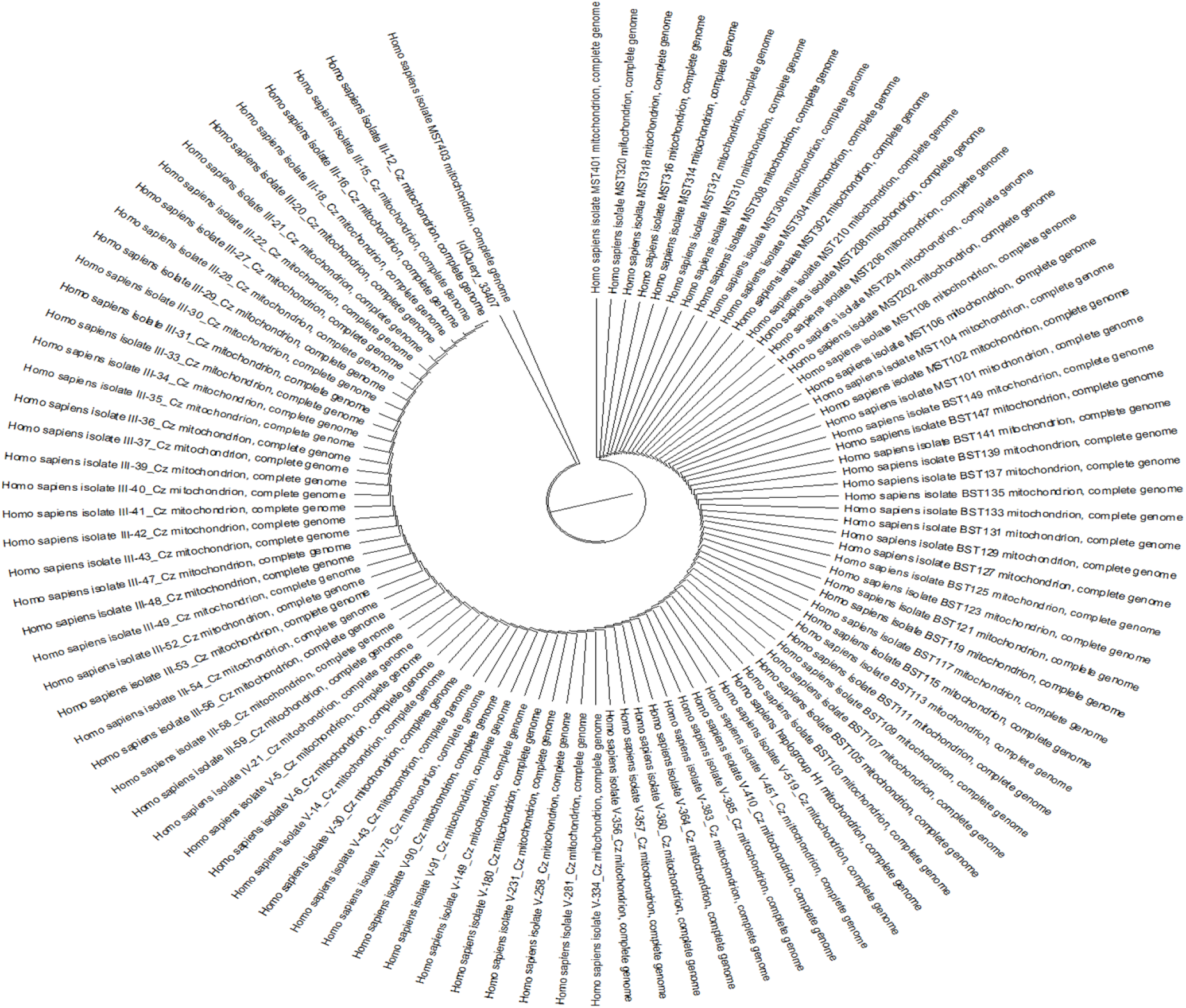
Showing the phylogenetic tree of the subgroups.

**Table 1.** Predicted Haplotype distribution and variant sequences of the Ameru mtDNA.

| Haplogroup | Proposed Origin | Ameru (%) | Chuka | Igembe | Igoji | Imenti | Muthambi | Mwimbi | Tigania | Tharaka |
| --- | --- | --- | --- | --- | --- | --- | --- | --- | --- | --- |
| K | Near East/ Jewish /Druze population | 4 |  |  |  | 17 | 8 |  | 12 |  |
| L0 | South East Africa | 20 | 21 | 13 | 21 | 33 | 33 | 14 | 12 | 13 |
| L1 | Central Africa/ West Africa | 3 | 7 | 7 |  |  |  | 7 |  | 7 |
| L2 | West and Central Africa | 6 | 7 | 13 | 7 |  | 8 |  |  | 6 |
| L3 | East Africa | 40 | 50 | 40 | 21 | 25 | 17 | 57 | 38 | 63 |
| L4 | Ethiopia, Somalia, and Kenya, Tanzania hunter and gatherer | 15 |  | 13 | 36 | 17 | 25 | 7 | 25 | 6 |
| L5 | E. Africa, great rift valley | 2 |  |  | 7 |  |  |  |  | 6 |
| M | Arab peninsula/ Near east | 5 | 7 |  | 7 |  | 8 | 7 |  |  |
| N | near east | 1 |  |  |  | 8 |  |  |  |  |
| P | South East Asia | 1 |  |  |  |  |  | 7 |  |  |
| R | South East Asia | 1 |  |  |  |  |  |  | 13 |  |
| V | South west europe | 1 | 7 |  |  |  |  |  |  |  |

It is important to note that haplogroup L_0_ (at 20%) haplotype is associated with the people living in Southern Africa. L_1_ haplogroup (at 3%), dominantly found among the Bantus of West Africa, Central Africa, and parts of Eastern Africa (Tanzania). L_3_(40%) is found in East Africa but is associated with Southern dispersal events to Asia. L_4_ (15%), whose origin is East Africa, especially those in Ethiopia, Somalia, and Kenya. Except for the Tharaka people, all the other subtribes had mtDNA haplotypes from outside of Africa. This shows a homogeneous group with no admixture genotype in the population.

According to the haplogroup, L1 haplogroup (2.9%) was found among Igembe (7%), Tharaka (6%), and Chuka (7%) and is common in West, Central, and parts of East Africa. L2 haplogroup (6.7%) was present in all subgroups except Imenti and Tigania, indicating West and Central African maternal ancestry. L1 and L2 haplotypes indicate that most Ameru subgroups share partial maternal ancestry from West and Central Africa, while Imenti and Tigania have different maternal lineages. Haplogroup L3, was most common among Tharaka (63%), Mwimbi (57%), and Chuka (50%) but absent in Imenti. Imenti lack L1 haplogroups and also do not carry L2, which is present in most other Ameru subgroups. This absence suggests that the Imenti have distinct maternal ancestry and did not share the West or Central African maternal lineages

**Table 2.** Showing subgroup haplotypes of the Ameru population.

| Haplogroup | Proposed Origin | Ameru (%) | Chuka | Igembe | Igoji | Imenti | Muthambi | Mwimbi | Tigania | Tharaka |
| --- | --- | --- | --- | --- | --- | --- | --- | --- | --- | --- |
| L1 | Central Africa/<br>West Africa | 3 | 7 | 7 |  |  |  | 7 |  | 7 |
| L2 | West and Central<br>Africa | 6 | 7 | 13 | 7 |  | 8 |  |  | 6 |
| L3 | East Africa | 40 | 50 | 40 | 21 | 25 | 17 | 57 | 38 | 63 |
| L4 | Ethiopia, Somalia,<br>and Kenya,<br>Tanzania hunter<br>and gatherer | 15 |  | 13 | 36 | 17 | 25 | 7 | 25 | 6 |

Haplotype L5 was identified mainly among the Igembe (7%), Igoji (7%), and Tharaka (6%) subgroups that points at Tanzania as their migration route. Haplogroup N was observed only among the Imenti subgroup (8%), suggesting possible Asian-related maternal ancestry. Haplogroup R (1%), found only in the Tigania subgroup, indicates early interaction with Asian populations or shared ancestry with the Horn of Africa.

The existential myth of the Ameru population having brotherhood with the Israelites was partly evident among other groups in the population but and not all. This was supported by the finding that the haplogroup K, originating mainly from the Middle East, was found exclusively among the Imenti (17%), Tigania (13%), and Muthambi (7%), indicating Middle Eastern ancestry. Absence of haplogroup K in other Ameru subgroups (Igembe, Igoji, Tharaka, Chuka, Mwimbi) indicates that Middle Eastern origin is inapplicable to them meaning that they have no sibling association with the Israelites. From the cladogram we found five groups that clustered together. Group 1: comprised of Muthambi, Mwimbi, Igembe set cluster together to infer close genetic relatedness. Moreover, Muthambi is genetically distant and diverse from the rest. However, Muthambi and Mwimbi are more closely (0.00066 units distance apart) related genetically. Group 2: comprised of Muthambi, Tigania, Mwimbi and Tharaka. Group 3: comprised of Igembe and Igoji while the Group 4: comprised of Chuka and Igoji. Then Group 5: comprised of the Imenti and Tharaka. Their diversity distance for this data set had bootstrap values lower than 0.2- -0.01 %. This means that the sequences were so similar such that, when a comparator on ClustalO to generate a tree was used, all the sequences collapse into one line clade (Fig. 2)

## Discussion

Based on mtDNA sequences there are two major migratory routes from Africa (Maca-Meyer et al., 2001; Gaitien et al., 2013). The southern route representing the haplogroup M expansion can be traced from Ethiopia through the Arabian Peninsula to India and Eastern Asia. However, the M haplogroup diversity is greater in India (Kivisild et al., 1999) than in Ethiopia (Quintarna-Murci et al., 1999). The northern route split into three main clusters. The first cluster comprising of the haplogroups W, I and N1b are found in Europe, The Middle East and Caucasia and also in Egypt and Arabian Peninsula. The next group divided into haplogroup X and A, common in Europe and Asia respectively (Guha et al., 2013; Bandelt et al., 2000). The third cluster subdivided into four lineages of which the first one gave rise to haplotype B found in Japan, East Asia, and Southern Pacific Archipelago, the second formed haplogroups J and T, whereas H and V, belong to the third cluster, their derivatives being found in Europe, North Africa and Central Asia (Marrerro et al., 2016; Maca-Meyer 2001). The fourth lineage is U and the highest frequencies of its sub-haplogroups are found in India (U2, U7), North Africa (U6, U3) and in Europe (U5; Witas and Zawicki, 2004). According to Richards et al. (1998), the major European mtDNA lineages are U5, H, I, J, K, T, V, W, and X. Haplogroup J encompassing about 16% of European mtDNA content, is probably the only one imported to Europe by the neolithic farmers. Recent studies indicate an early invasion of a single, ancestral lineage of Asian origin in America (Bonatto and Salzano, 1997; Silva et al., 2002). The four most common American haplogroups – A, B, C, and D, although old, have similar nucleotide polymorphism, suggesting their common origin (Silva et al., 2002).

Haplotype diversity is a measure of the uniqueness of a particular haplotype within a population, reflecting the genetic variation within a group. Within these groups are ancestral haplotypes that refers to the haplotype of a most recent common ancestor, which can be deduced by comparing haplotypes of descendants and eliminating mutations a process referred to as triangulation, that can be used to identify the genetic makeup of a shared ancestor (Allard et al., 2002; Miller et al., 2001). In this regard, the use of mtDNA analysis has always pointed to the Amerus’ as having a close genetic relationship with the Adean Ameridians population with their haplotypes having a distinct combination of genetic markers inherited together on a chromosome, specifically within the Amerindian population that provide a genetic history, migration patterns, and relationships between different Amerindian groups (Achieli et al., 2005; Pereira et al., 2010). However, the Ameridian population display a geographical pattern in the frequencies of different mtDNA haplotypes (A, C, and D). The frequencies showed geographical variation, with the highest frequencies in the Andes and North Amazon regions an indication of the Amerus’ originating from the Amazon. (Macaulay et al., 1999) The emergence of haplogroup L1-6 at a later time, possibly in a population in eastern Africa is also reported. In these, L1, L5 and L2 are reported to be out of Africa though found in Africa indicating a genetic drift into Africa. Malyarchuk et al., 2008; Salas et al., 2004; 2005; Veeranch et al., 2020; Stettova et al., 2011) The “out of Africa” hypothesis proposes that a small group of Homo sapiens left Africa 80,000 years ago, spreading the mitochondrial haplotype L3 throughout the Earth outside Africa. L3 that emerged much later was found to be closely associated with the out-of-Africa event but it may have arisen either in East Africa or in Asia. Within L3 came L6 and L4 as sister clades that are only limited to East Africa and did not participate in the out-of-Africa migration. Haplotype L_4b2a2_ (Kenya), L_2a1a2_ (Tanzania), L_0a_ (Tanzania), L_4b2a2a_ (Tanzania) and L_2a1_ (Tanzania) are associated with the east Africa region. However, L0d2c1, L11b2b1b and L0d1a1a are found in South Africa. (Quintana-Murci et al., 1999; Wallace et al., 1999) It is therefore inferred that an earlier wave of expansion of *Homo sapiens* left Africa and left genetic traces by interbreeding with Neanderthals before disappearing. Ancient individuals from Ethiopia (∼4500 BP), Kenya (∼400BP), Tanzania (both ∼ 1400BP) and Malawi (∼ 5000Years) show increased affinity to South African’s (Mendell et al., 2010). The L haplogroups are reported to be predominantly allover sub-Saharan Africa viz:- North Africa {i.e. central, eastern and west African populations have experienced episodes of heavy exchange and gene flow (e.g. expansions within haplogroup L3}, Sudan, Ethiopia, West Africa, East Africa {Kenya, Uganda, Tanzania}, Southern Africans, Southeast Africa {Mozambique), Native Southern African {!Xung, !Kung and Khwe khoisans), Mbenga Pygmies {Baka, Bi-Aka and Ba-Kola}, Ba-Mbuti, Pygmies, Hadza/Sandawe). However, they seem to radiate from Africa through North Africa, Arabia Peninsulas, Middle East, and West Asia, {most notably in Yemen} (Alvarez-Iglesias et al., 2009) . While in the Middle East, the haplotype is found in Bedouins from Israel, Palestinians, Jordaninans, Iraqis, Syrians, Hazara of Afghanistan, Saudi Arabians, Lebanese, Druzes from Israel, Kurds and Turks. In addition, America, and Europe also has the L haplotype.

In our study, 75% of the Ameru’s had mtDNA Haplogroup L. Geographically, this place their origin to be from Africa then to the Middle East through North Africa, Arabia Peninsula to West Asia. The haplotype also undergone extensive back-flow from Eurasian populations (via North Africa or Arabia/East Africa) long after the initial dispersal out of Africa, which has reduced their differentiation from non-Africans.

Haplotype L is subdivided into haplotype L_1-6_ of which L_1_, L_5_, and L_2_ are found out of Africa and are associated with haplotype A, C, and D which is found in Adean Ameridian group geographically found in Southern America {from North to South of the Amazon River} population(Maria et al., 2012; Watson et al., 1997). L1 is associated with Africa (Congo, Gabon, Cameroon), while L_5_ (L_1e_) is associated with Mbuti Pygmies in East and Central Africa. L_2_ (L_1_, L_4_, L_5_) is found in East and West Africa then migrate south Eastern (Bantu migration) to Central sub-Saharan Africa at the expense of L_0_, L_1_, L_5_ (Fernades et al., 2012; Chen et al., 2000). L_3_ is associated with Southern dispersal event out of Africa migration. It is possibly in East Africa (mtDNA) that all of non-Africans is derived from and is divided into two main lineages of M and N (. Haplotype N is proposed to originate from East Africa and Asia with L3 as an ancestor (Winter et al., 2010). People carrying mtDNA N lineages could have reached Australia following a northern route through Asia.The northern route of migration via North Africa and the Nile valley into the Levant with subsequent dispersal into both Europe and Asia. M haplotype is thought to have originated from East Africa, Southwest Asia, Southeast Asia or South Asia (Winters et al., 2014). It descends from L3 haplotype that is associated with Southern dispersal event out of Africa. Through this route humans left Africa by crossing the Bab-el-Mandeb strait at the mouth of the Red Sea and then rapidly migrated along the South Asia coastline to Australia/Melancesia. The migration was across the Red Sea and along the east coast of Arabia. This group spread around the coast of Arabia and Persia until they reached India. From Arabia to India the proportion of haplogroup M increased eastwards (from India along the coasts of Thailand and Indonesia all the way to Papua New Guinea. The ancestral L_3_ lineages were then lost by genetic drift in Asia as they are infrequent outside Africa. M macrohaplogroup is then hypothesised to originate from Asia (specifically in the Indian subcontinent, China, Japan and Korea). India has several M lineages that emerged directly from the root of haplogroup M. Only two subclades of haplogroup M, M_1_ and M_23_, are found in Africa, whereas numerous subclades are found outside Africa. M_1_ in Africa is predominantly North African/supra-equatorial and is largely confined to Afro-Asiatic speakers, which is inconsistent with the Sub-Saharan distribution of sub-clades of haplogroups L_3_ and L_2_ that have similar time depths. One of the basal lineages of M_1_ lineages has been found in Northwest Africa and in the Near East but is absent in East Africa. It is relatively common in the Mediterranean, Middle East and Central Asia. The origins of haplogroup M further complicated by an early back-migration (from Asia to Africa) of bearers of M_1_. The haplotype M, also referred to as M_23_, is present in Madagascar. Thus, the presence of M_1_ in Africa is the result of a back-migration from Asia through M_23_ which occurred sometime after the Out of Africa migration. This far, it is indicative that the Ameru population like any other Bantu speaking group migrated through the horn of Africa to the Middle East, Arabia Peninsula to West Asia then back to Africa through Madagascar. This is the hypothetically migratory route of the Ameru.

Haplogroup L0 ((L_0a’b’f_)) emerges from the basal haplogroup L1-6. L0 is the most divergent clade in the maternal line of descent. The haplotype is associated with the people living in Southern Africa and Namibia. The haplotype subclades are L_0d_ and L_0k_. Haplogroup L_0_ consist of four main branches (L_0d_, L_0k_, L_0a_, L_0f_). All of them were originally classified into haplogroup L_1_ as L_1d_, L_1k_, L_1a_ and L_1f_. Haplotype L_0d_ and L_0K_ are associated with the Khoisan of South Africa but L_0d_ has also been detected among the Sandawe people of Tanzania, which suggests an ancient connection between the Khoisan and East African speakers. Haplogroup L_0f_ is present in relatively small frequencies in Tanzania among the Sandawe people who are known to be older than the found in Ethiopia. Inferably, haplotype L_0a_ is majorly found in South Africa and Southern East Africa (Tanzania), making the route form M_1_ (M_23_) in Madagascar to South Africa, Namibia, Botswana then Mozambique and then to Tanzania (Sun et al., 2006; Fenardes et al., 2012). The haplotype is equally found in Yemen confirming out of Africa migration. Haplotype L0 (L_0a_’_b_’_f_) may have an eastern African origin (Shi et al., 2010). Based on our study analysis 20% of Ameru blood samples were of L_0_ haplotype. This shows a relationship with the Khoisan and East African speakers particularly in regions like Ethiopia, Kenya, and Tanzania. This implies that they made their route back through Madagascar as M_1_ (M_23_) to South Africa, Namibia, Botswana, Mozambique, Tanzania and probably to the coastal Kenya (Soares et al., 2012; Macaulay et al., 2005; Palanichamy et al., 2004; Torroni et al., 2005). About 2.9% and 6.7% were respectively members of L_1_ haplogroup, predominantly found among the Bantus and Semi-Bantus in West Africa (Congo, Gabon, Cameroon), Central Africa, and parts of Eastern Africa (Tanzania). 40% of the Ameru haplogroup L_3_ was found in East Africa and is associated with Southern dispersal event to Asia where the haplotype disappeared due to genetic drift and was taken over by haplotype M (found in the Philippines, Adaman Sea Island, New Guinea, Australia and Malaysia)(Behar et al., 2012). 15.2% of the analyzed blood samples were of the haplogroup L_4_ (L_4b2a2_), whose origin is East Africa especially those in Ethiopia, Somalia, and Kenya but is also present in some regions of South Africa and the Horn of Africa but did not participate in the out of Africa migration hence no relationship with the L_3_ haplotype. Haplotype L_5_ was found in 1.9% of participants. The haplotype is primarily found in East and Central African populations, including among the Hadza people of Tanzania, and the Mbuti pygmies (Huang et al., 2011).

In conclusion, Ameru like the other Bantu speakers, originated from Africa and migrated through the horn of Africa to the Middle East, Arabia Peninsula to West Asia then back to Africa through Madagascar, South Africa, Namibia, Botswana, Mozambique, and Tanzania and then Kenya. Their northern route of migration was via North Africa and the Nile valley into the Levant with subsequent dispersal into both Europe and Asia (Michael C Campbell and Sarah A Tishoff, 2010). Through this route humans left Africa by crossing the Bab-el-Mandeb strait at the mouth of the Red Sea and then rapidly migrated along the South Asia coastline to Australia/Melancesia. The migration was across the Red Sea and along the east coast of Arabia. The spread was around the coast of Arabia and Persia until they reached India. From India the group migrated along the coasts of Thailand and Indonesia all the way to Papua New Guinea, Philippines, Adaman Sea Island, Australia and Malaysia. From Asia, they took a migratory route form to Madagascar then to South Africa, Namibia, Botswana, Mozambique and then to Tanzania (Phillip et al., 2007; Metspalu et al., 2006; Macaulay et al.,2005). Upon reaching East Africa, there was a group that migrated to West Africa (Congo, Gabon, Cameroon), Central Africa, and parts of Eastern Africa (Tanzania, Ethiopia, Somali and Kenya).

## Conclusion

In conclusion, the Ameru population exhibits diverse maternal ancestry, reflecting contributions from multiple African and non-African lineages: L0–L4 haplogroups indicate predominant East, Central, and West African maternal origins, with subgroups showing variation in haplotype frequencies (e.g., L1 and L2 in Igembe, Tharaka, Chuka; L3 in Tharaka, Mwimbi, Chuka; L4 across all subgroups). Subgroup differences suggest that certain communities, particularly Imenti, have distinct maternal lineages, with less contribution from L1, L2, and L3 but potential links to Afro-Asiatic groups via L4 (found in the Middle East). Non-African haplogroups (N and R) point to historical interactions or shared ancestry with populations in Eurasia and the Horn of Africa, primarily in Tigania and Imenti. Overall: The Ameru maternal gene pool is heterogeneous, shaped by multiple migration routes and interactions across East Africa and beyond, with subgroup-specific maternal histories. Therefore, Ameru like the other Bantu speakers, originated from Africa and migrated through the horn of Africa to the Middle East, Arabia Peninsula to West Asia then back to Africa through Madagascar, South Africa, Namibia, Botswana, Mozambique, and Tanzania and then to Kenya. Their northern route of migration was via North Africa and the Nile valley into the Levant with subsequent dispersal into both Europe and Asia. Through this route humans left Africa by crossing the Bab-el-Mandeb strait at the mouth of the Red Sea and then rapidly migrated along the South Asia coastline to Australia/Melancesia which supports our folklore findings of the Ameru migratory route. The migration was across the Red Sea and along the east coast of Arabia. The spread was around the coast of Arabia and Persia until they reached India. From India the group migrated along the coasts of Thailand and Indonesia all the way to Papua New Guinea, Philippines, Adaman Sea Island (in Indian Ocean near Indonesia and Thailand), Australia and Malaysia (Asia). From Asia, they took a migratory route back to Madagascar then to South Africa, Namibia, Botswana, Mozambique and then to Tanzania. Upon reaching East Africa, there was a group that migrated to West Africa (Congo, Gabon, Cameroon), Central Africa, and parts of Eastern Africa (Tanzania, Ethiopia, Somali and Kenya).

## Acknowledgement

This study was sponsored by Mr. Raphael Anampu who had the quest to know about the Origin and Migration patterns of the Ameru community in Kenya. We thank the Ameru community who presented themselves as study participants.

